# Human Ageing Genomic Resources: updates on key databases in ageing research

**DOI:** 10.1101/2023.08.30.555622

**Authors:** João Pedro de Magalhães, Zoya Abidi, Gabriel Arantes dos Santos, Roberto A. Avelar, Diogo Barardo, Kasit Chatsirisupachai, Peter Clark, Evandro A. De-Souza, Emily J. Johnson, Inês Lopes, Guy Novoa, Ludovic Senez, Angelo Talay, Daniel Thornton, Paul Ka Po To

## Abstract

Ageing is a complex and multifactorial process. For two decades, the Human Ageing Genomic Resources (HAGR) have aided researchers in the study of various aspects of ageing and its manipulation. Here we present the key features and recent enhancements of these resources, focusing on its six main databases. One database, GenAge, focuses on genes related to ageing, featuring 307 genes linked to human ageing and 2205 genes associated with longevity and ageing in model organisms. AnAge focuses on ageing, longevity, and life-history across animal species, containing data on 4645 species. DrugAge includes information about 1097 longevity drugs and compounds in model organisms such as mice, rats, flies, worms, and yeast. GenDR provides a list of 214 genes associated with the life-extending benefits of dietary restriction in model organisms. CellAge contains a catalogue of 866 genes associated with cellular senescence. The LongevityMap serves as a repository for genetic variants associated with human longevity, encompassing 3144 variants pertaining to 884 genes. Additionally, HAGR provides various tools as well as gene expression signatures of ageing, dietary restriction, and replicative senescence based on meta-analyses. Our databases are integrated, regularly updated, and manually curated by experts. HAGR is freely available online (https://genomics.senescence.info/).

## Introduction

Ageing is one of the most complex biological processes, yet despite extensive studies, its underlying mechanisms remain to be elucidated (1,2). Furthermore, ageing is a major risk factor for mortality and several diseases, leading researchers from various fields to study this process (3). The Human Ageing Genomic Resources (HAGR) is an intuitive and powerful collection of online tools and databases that have greatly assisted scientists in addressing this complex problem.

HAGR first became publicly available online in 2004 and has been growing dynamically since, in parallel with the major growth and development of ageing-related research (4). Our resources include six main databases related to different aspects of ageing research (Figure 1), alongside other supplementary projects and general information on ageing biology. Given the impact of genetics on ageing, HAGR places a strong emphasis on genetic and genomics.

**Figure 1:**
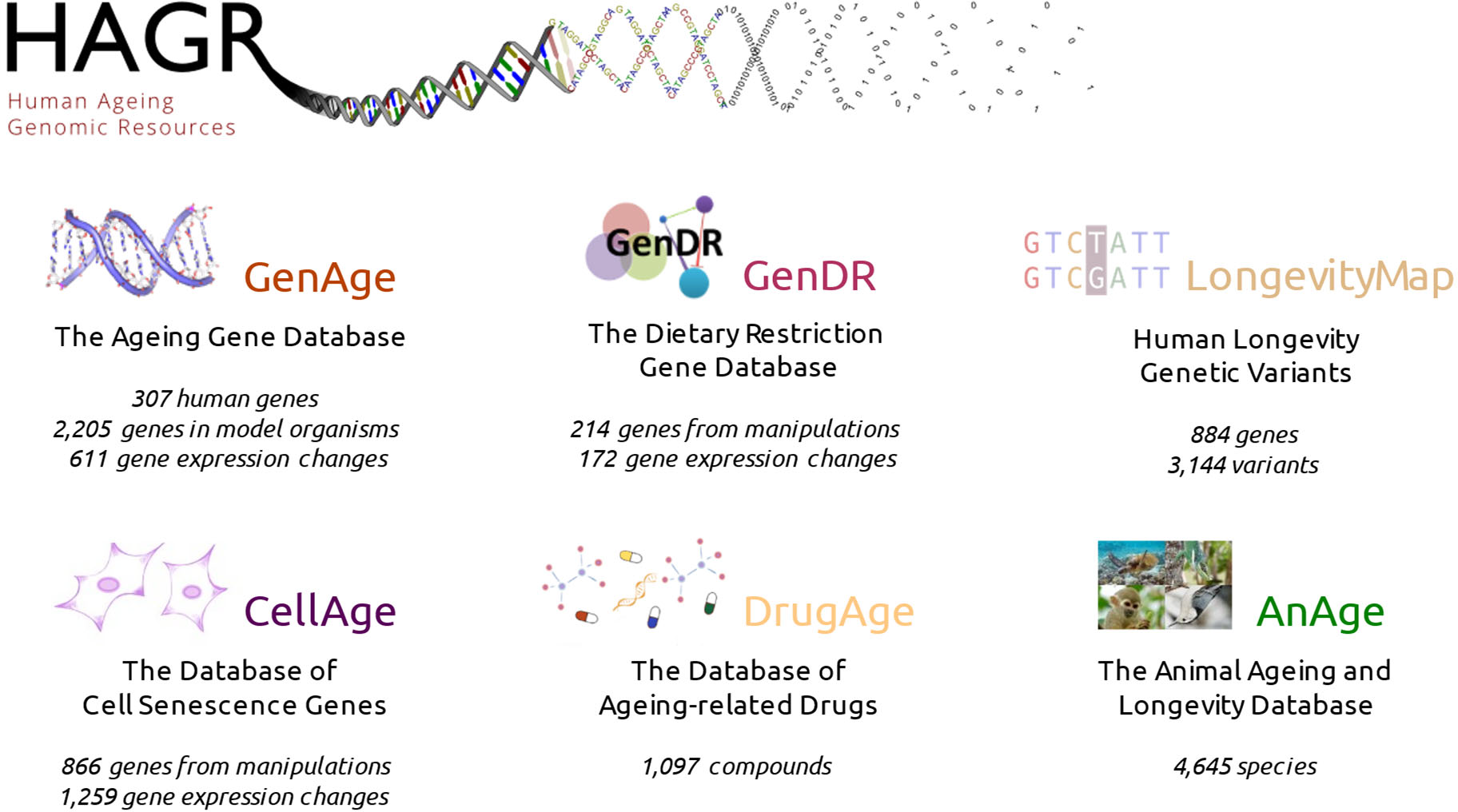
Overview of the main databases in the Human Ageing Genomic Resources (HAGR).

In this paper, we provide an overview of the current version of HAGR, summarizing each of its databases and tools and highlighting updates made since its previous release (5). Our goal is to provide an up-to-date guide to one of the leading and most frequently accessed online platforms used in biogerontology. HAGR is freely available at https://genomics.senescence.info/, with no registration required.

## Online Resources and Databases

### GenAge: The database of ageing-related genes

GenAge (https://genomics.senescence.info/genes/) is our benchmark database focused on genes associated with the ageing process, the so-called “gerontome”. It is in turn divided into two core databases: i) GenAge – Human; and ii) GenAge – Model Organisms. Both of these have been manually curated from the scientific literature. Detailed information about GenAge is available in earlier publications (6), what follows is a brief description.

GenAge – Human – is a repository of genes associated with human ageing, or at least genes that may significantly impact the human ageing phenotype and processes. It is important to note that our focus within HAGR is on biological ageing, not just age-related diseases. Because longevity is influenced by multiple factors beyond ageing, such as accidents, we focus on genes that potentially regulate the ageing process as a whole – or that at least influence various aspects of the ageing phenotype – rather than solely those affecting longevity (refer to the LongevityMap below for longevity-related genes). Each human gene entry was thus selected following a review of the literature and considering that genes can be associated with ageing based on different types of studies and evidence. These studies include human genetic association studies, genetic manipulations in model organisms, and in vitro studies. Consequently, genes are classified into nine categories corresponding to their level of evidence (ranging from “indirect/inconclusive evidence linking the gene product to ageing” to “evidence directly linking the gene product to ageing in humans”).

Genes commonly differentially expressed during mammalian ageing are also available to researchers in GenAge. A recent meta-analysis by our lab revealed global and tissue-specific gene expression changes during human ageing, with significant overlaps with both GenAge Human and with the LongevityMap (7). More specifically, we identified 449 upregulated and 162 downregulated genes with age across all tissues. This is a substantially larger mammalian ageing signature than previously, which consisted of 56 upregulated and 17 downregulated genes (8), possibly due to the larger number of studies now available.

GenAge – Model Organisms – comprises a list of genes associated with longevity or ageing in model organisms based on genetic manipulation experiments curated from the literature. To be clear, only genes that, when genetically manipulated, have a significant impact on ageing and/or longevity are included. Genes are classified into three categories: “pro-longevity”, “anti-longevity”, and “necessary for fitness”. Genes with conflicting results or insufficient data are unannotated and categorized as “unclear”. Information from model organisms is also leveraged to infer possible genes associated with human ageing in the aforementioned human dataset. The species distribution in GenAge is presented in Table 1 and includes the major traditional biomedical models such as mice, flies, worms, and yeast.

**Table 1.** Species in the GenAge database.

| **Database** | **Species** | **Number of genes** |
| --- | --- | --- |
| Human genes | *Homo sapiens* | 307 |
| Model organism genes | *Mus musculus* | 136 |
|  | *Caenorhabditis elegans* | 889 |
|  | *Drosophila melanogaster* | 202 |
|  | *Saccharomyces cerevisiae* | 911 |
|  | *Caenorhabditis briggsae* | 1 |
|  | *Danio rerio* | 1 |
|  | *Mesocricetus auratus* | 1 |
|  | *Podospora anserina* | 3 |
|  | *Schizosaccharomyces pombe* | 61 |
|  | **Total for model organisms** | 2205 |

While the number of human ageing genes has only modestly changed over the years (5), existing entries are further curated, resulting in the addition of dozens of new bibliographic references. Observations concerning many genes are also regularly updated to reflect new findings. In regard to model organisms, in addition to including almost 100 new genes since the last update, we also added another five additional species, most notably *S. pombe*, an essential unicellular organism in ageing research – now featuring 61 entries (9,10).

### AnAge : The database of animal ageing and longevity

AnAge (https://genomics.senescence.info/species/) is a curated database of ageing, longevity and life-history traits. Its primary aim is to support studies involving comparative ageing biology while also being of value to fields such as ecology and conservation.

For a comprehensive description of AnAge, please refer to an earlier publication (11). In brief, this database presents entries with a multitude of data, including data on maximum lifespan, metabolism, taxonomy, and additional life-history data, alongside relevant ageing phenotypes and observations. AnAge incorporates over 1400 articles and also highlights a list of 9 species exhibiting negligible senescence.

Although primarily focused on animal biology, AnAge also includes entries about plants (n=5) and fungi (n=4). Regarding animals, nine phyla are represented, with approximately 98% of the entries in Chordata, divided into 14 classes. The most well-represented classes are *Aves, Mammalia, Teleostei*, and *Reptilia*, respectively (Table 2). AnAge is of great value for ageing research as offers information on the wide range of lifespans across taxa (Figure 2), facilitating a variety of comparative studies in biogerontology.

**Table 2.** Taxonomy of species in the AnAge database.

| **Kingdom *Animalia*** |  |  |
| --- | --- | --- |
| **Phylum** | **Number of species** | **Average longevity (years)** |
| *Annelida* | 3 | 283.3 |
| *Arthropoda* | 16 | 11.6 |
| *Chordata* | 4557 | 19 |
| *Cnidaria* | 2 | NA |
| *Echinodermata* | 2 | 125 |
| *Mollusca* | 49 | 36 |
| *Nematoda* | 2 | 0.6 |
| *Platyhelminthes* | 3 | 2.3 |
| *Porifera* | 2 | 8275 |
| **Phylum *Chordata*** |  |  |
| **Class** | **Number of species** | **Average longevity (years)** |
| *Amphibia* | 181 | 14.9 |
| *Actinopterygii* | 4 | 27.4 |
| *Ascidiacea* | 1 | 2 |
| *Aves* | 1513 | 17.8 |
| *Cephalaspidomorphi* | 16 | 7.1 |
| *Chondrichthyes* | 117 | 25.5 |
| *Chondrostei* | 14 | 71.2 |
| *Cladistei* | 1 | 34 |
| *Coelacanthi* | 1 | 48 |
| *Dipnoi* | 3 | 37.1 |
| *Holostei* | 4 | 26 |
| *Mammalia* | 1349 | 19.8 |
| *Reptilia* | 547 | 22 |
| *Teleostei* | 806 | 16.9 |

**Figure 2:**
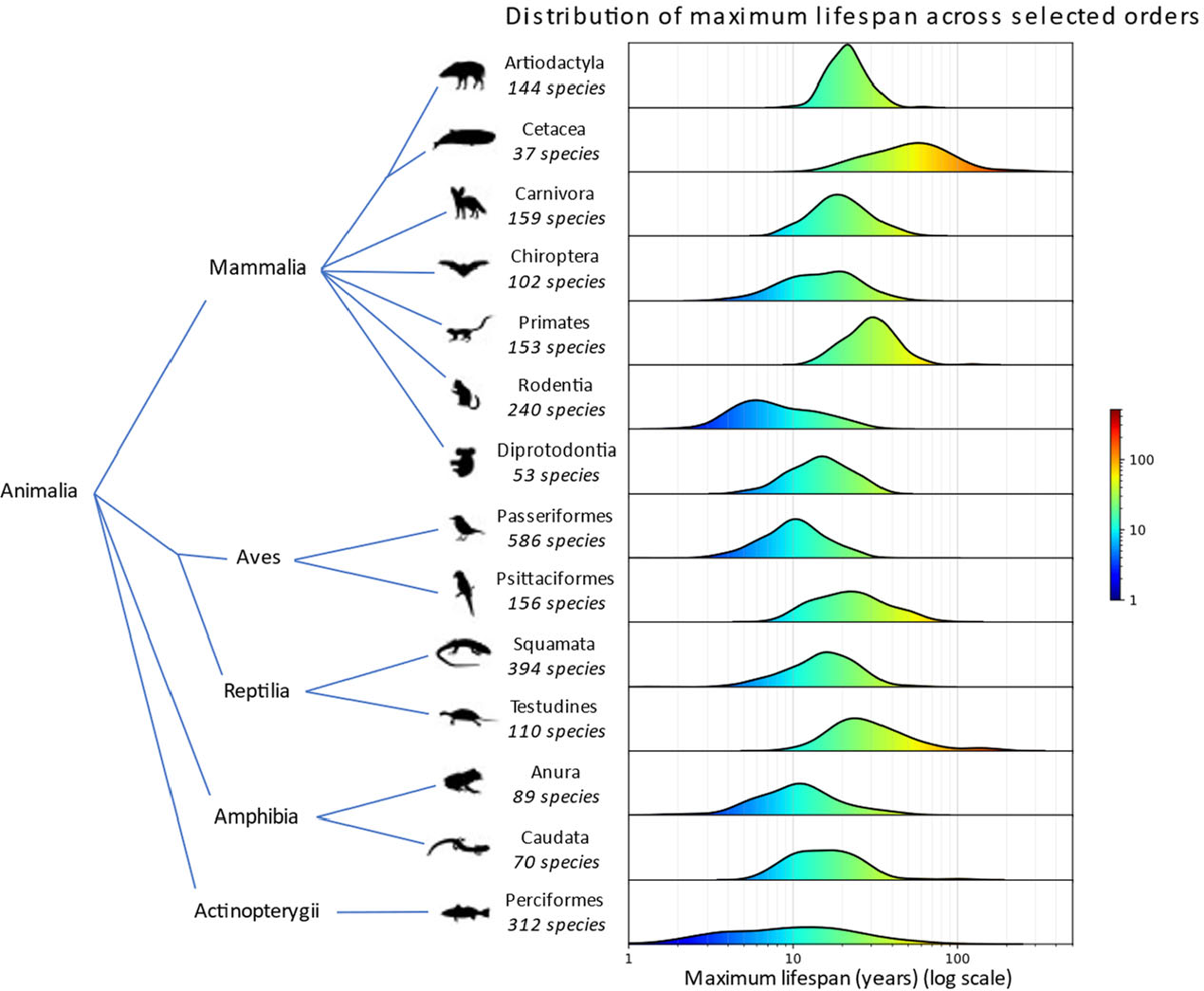
Distribution of maximum lifespan across selected orders using data from the AnAge database. Six mammalian orders, two bird, two reptile, two amphibian, and one fish order (each with over 40 species in the AnAge database) are included. Kernel density estimation was used for the distribution with default parameters. In the maximum lifespan distribution graph, the x-axis represents the maximum lifespan in years (log scale) while the y-axis indicates the number of species at each lifespan.

AnAge is one of the oldest and most frequently used HAGR resources and is arguably the benchmark animal longevity database worldwide due to its constant updating and manual curation. The current version comprises 4671 entries, encompassing 4645 species and 26 taxa.

### DrugAge: The database of anti-ageing drugs

Among HAGR’s recent additions, DrugAge (https://genomics.senescence.info/drugs/), consists of a manually curated compilation of drugs and compounds that extend longevity in model animals (12). Some compounds are listed multiple times because they have been tested across various species and doses, enabling more comprehensive assessments of their impact on longevity, as shown previously using DrugAge (13). Given our focus on ageing, compounds from studies involving disease-prone animals or harmful conditions are not included.

Presently, DrugAge features 1097 drugs or compounds evaluated in over 3200 experiments across 37 species, supported by a total of 656 references. Figure 3 illustrates the growth trajectory of HAGR databases over the years, where we can see a major improvement over the previous versions that is more marked for DrugAge. Indeed, longevity pharmacology has been exploding, and the growth in longevity drugs has outpaced the growth of longevity genes (14).

**Figure 3:**
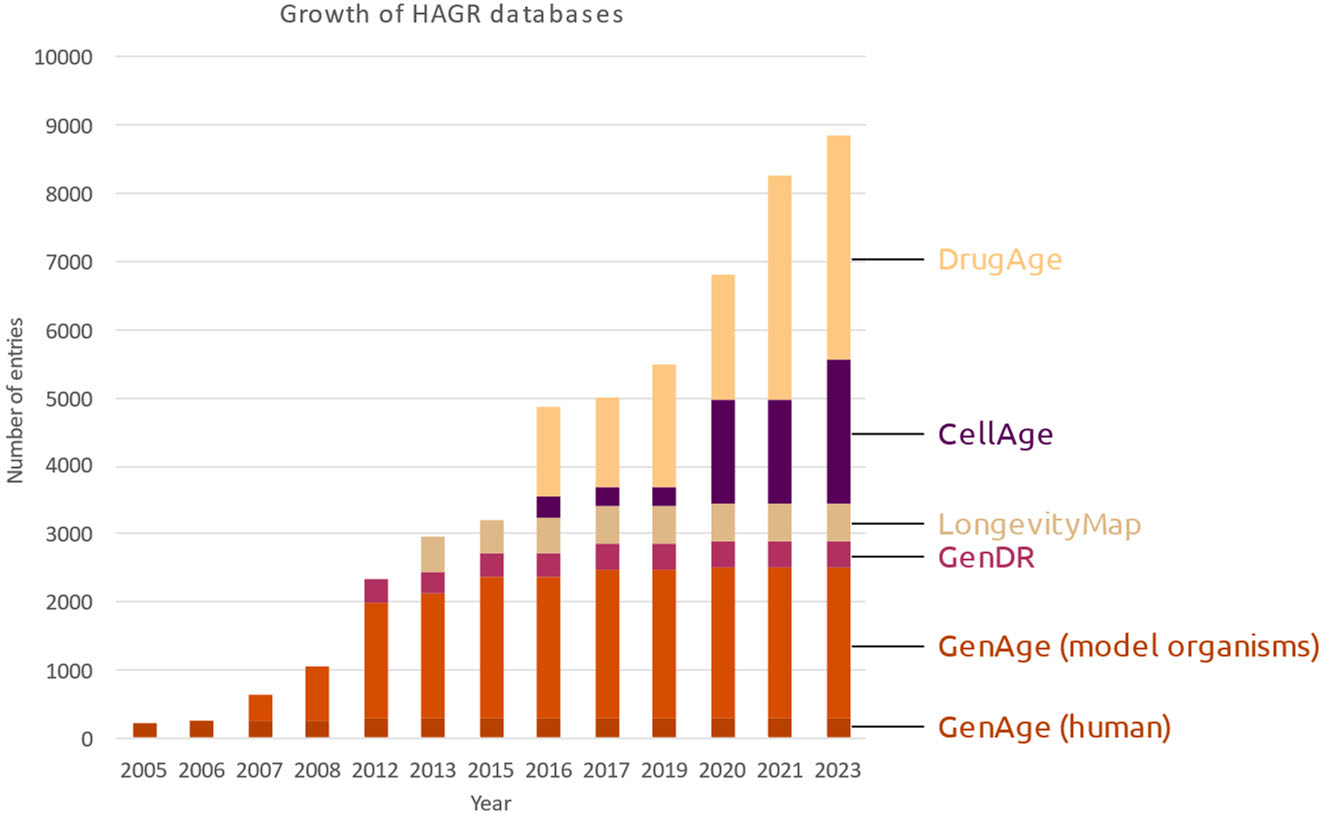
Time-series growth of the Human Ageing Genomic Resources (HAGR) databases. The figure illustrates the number of entries across different HAGR databases over the years. Gene expression signatures are not included.

The recent surge of interest in anti-ageing drugs within health research and the pharmaceutical/biotechnology sectors underscores the significance of a scientifically reliable resource like DrugAge (14,15). Despite its relatively recent development, our database has established itself as a leading information source in geroscience and is, after GenAge and AnAge, the most widely accessed database within HAGR (Figure 4).

**Figure 4:**
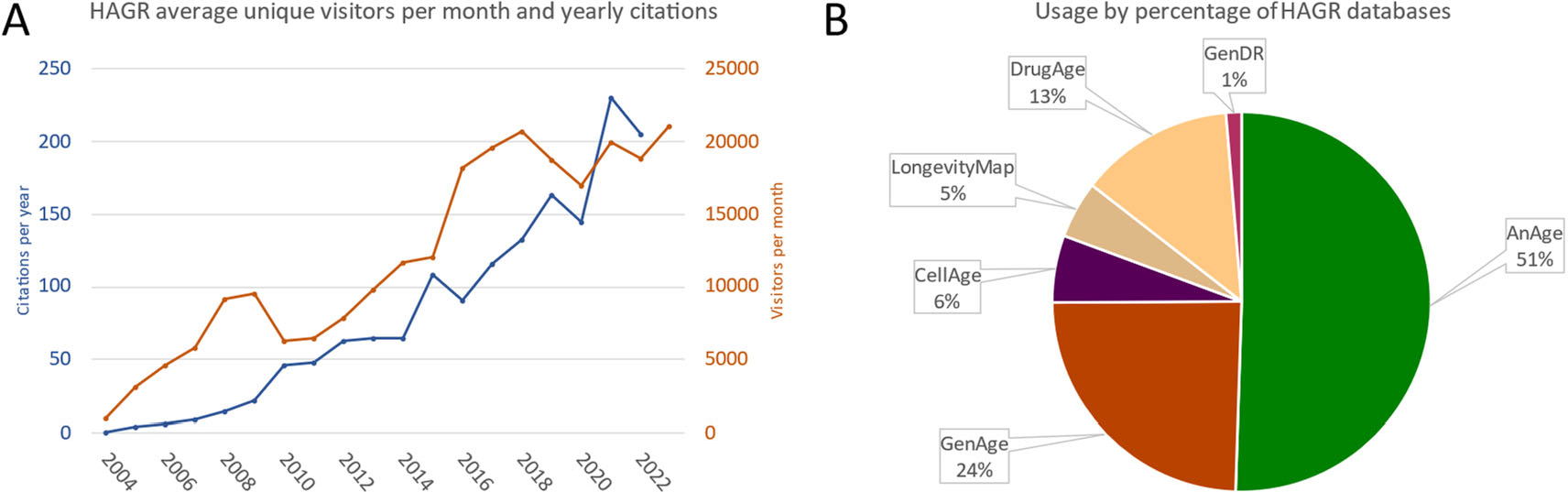
Human Ageing Genomic Resources (HAGR) visitors, citations, and database statistics. (A) HAGR unique visitors per month and citations per year of HAGR’s papers. (B) Usage by percentage of the different HAGR databases in 2022.

### GenDR: The database of dietary restriction genes

Dietary restriction (DR) is a widely studied anti-ageing intervention, yet its underlying mechanisms remains poorly understood (16-18). Recognizing the genetic component of ageing, our GenDR database (https://genomics.senescence.info/diet/) compiles genes associated with DR to aid research and advance our understanding of the genetic and molecular mechanisms of DR-induced life-extension.

Further details about GenDR are available in a previous publication (19), but briefly this database includes two datasets: i) genes inferred from genetic manipulation experiments in model organisms to regulate the life-extending benefits of DR; and ii) mammalian genes whose expression is robustly altered due to DR derived from a meta-analysis (20). In total, GenDR comprises five model organisms and entails 214 genes inferred from genetic manipulations, alongside 172 genes derived from gene expression changes. To our knowledge, it remains the only database of genetic alterations associated with DR.

### CellAge: The database of cell senescence genes

Cellular senescence is triggered by various stressors like telomere attrition during replication, the dysregulation of onco- and tumour-suppressor genes, and cellular or DNA damage from sources such as hydrogen peroxide and irradiation (21,22). Senescent cells undergo proliferative arrest and secrete a mix of proinflammatory factors known as the senescence-associated secretory phenotype (SASP) (23,24). As senescent cells continually produce proinflammatory factors they may contribute to inflammageing and hinder tissue repair and renewal (25). Cellular senescence has been linked to various ageing-related diseases including cancer, Alzheimer’s disease, osteoarthritis, and diabetes (26-29).

CellAge was compiled from a systematic search of the literature, and genes were included based on specific criteria, as described (30). Briefly, the CellAge database consists of genes inferred from genetic manipulations in vitro that induce (n=370, 42.7%) or inhibit (n=475, 54.8%) replicative (n=153), stress-induced (n=185), and oncogene-induced (n=238) cellular senescence (https://genomics.senescence.info/cells/). Some genes are involved in multiple classes of senescence. There are 21 genes that have an unclear effect on cellular senescence, inducing or inhibiting this process depending on experimental context. Additionally, there are 360 genes in CellAge where the mechanism by which they influence the senescence program is unclear.

The current version of the CellAge database contains 866 genes (31), a considerable increase from the 279 genes in the first build (30). Furthermore, we previously used ‘replicative senescence’ as the default annotation for CellAge genes when the literature did not directly specify how the gene was influencing the senescence phenotype. We have now added a fourth annotation, ‘unclear,’ alongside the ‘replicative,’ ‘stress-induced,’ and ‘oncogene-induced’ tags, in order to better represent our knowledge of how these genes contribute towards cellular senescence. Previous entries have been updated to reflect this new annotation where applicable.

Furthermore, CellAge includes a list of 1259 genes differentially expressed during replicative senescence (525 and 734 over- and underexpressed, respectively) derived from a meta-analysis of senescent cells compared to proliferating counterparts (32).

### LongevityMap: The database of genetic association studies of longevity

While human longevity derives from a complex interplay of factors, the heritability of human longevity has been estimated to be ∼25% (33). The LongevityMap (https://genomics.senescence.info/longevity/) was developed to assist in cataloguing the increasing volume of data arising from genetic-variant studies of human longevity (34).

Succinctly, all entries within the LongevityMap were curated from the literature, excluding studies in cohorts of unhealthy individuals at baseline. Details on study design are provided for each entry, including population details, sample sizes, and indications of statistical significance, alongside negative results. In total, our database encompasses over 500 entries, comprising 884 genes and 3144 variants. The list is derived from 270 individual studies and presents 275 statistically significant results.

### Additional datasets, tools, and features

In addition to our core databases, HAGR also integrates other relevant resources on ageing. Of note, the Digital Ageing Atlas (https://ageing-map.org/) serves as a platform for age-related changes that integrates age-related molecular, physiological, psychological, and pathological data to deliver an interactive database that centralizes human ageing-related changes (35). Despite its status as an external database to HAGR, we link to this portal from HAGR to provide additional ageing-related context to genes in HAGR. Likewise, our genomics resources include genome and transcriptome sequencing of the naked-mole rat (the longest-lived rodent) and the bowhead whale (the longest-lived mammal) (36,37). Moreover, HAGR encompasses projects on the relationship between ageing, cancer, and evolution. Beyond genes related to age-related diseases and transcriptional signatures of ageing, these projects host datasets that can be explored and downloaded. In addition, two bioinformatics tools for ageing research are featured in HAGR: Ageing Research Computational Tools (ARCT), a Perl toolkit, and an SPSS script to determine the demographic rate of aging for a given population, as described previously (38).

Furthermore, HAGR provides information and news about ageing biology. Our website includes links to social media (Twitter: https://twitter.com/AgingBiology and Facebook: https://www.facebook.com/BiologyAgingNews) with the latest news in the biology and genetics of ageing and includes an educational resource on ageing (https://www.senescence.info/). Lastly, we maintain WhosAge (http://whoswho.senescence.info/), a compilation featuring 340 individuals and 65 companies contributing to biogerontology.

#### Usage examples in ageing and longevity

With over 1,000 citations, HAGR has been invaluable to multiple studies in various and diverse research topics in the biology of ageing. Of note in recent years, AnAge was used in a study to identify species with remarkable longevity with evolutionary implications for lifespan (39). AnAge has also facilitated data gathering on species’ maximum lifespan and relative age comparisons (40,41), body size values in a study exploring evolutionary pathways to SARS-CoV-2 resistance (42), and age of female sexual maturity in a major research endeavour on somatic mutation rates across mammals (43). HAGR’s data contributed to accurate predictions of basal metabolic rate and organ weights (44).

To highlight other recent examples of the use of HAGR, Townes et al. used GenAge in their work to identify new potential genes associated with longevity in *C. elegans* and *S. cerevisiae* (45). Moreover, GenAge and DrugAge facilitated the annotation and curation of ageing-related genes/proteins which lead to the identification of potential drug targets (46). In collaboration with other scientists, our research group has used GenDR to aid in the application of machine learning methods to identify DR-associated features (47). A recent study used DrugAge as the primary source of information where they identified lifespan-extending compounds in diverse model organisms, providing novel insights on lifespan extension (48). A co-regulated network of senescence genes in human tissues was created using CellAge’s features (49). The CellAge database was also utilized in a study on microglial senescence, where human senescence signatures and senescence-associated genes were retrieved (50), and in another research endeavour where it assisted in identifying genes differentially expressed in aged stem cells (51). Further, the association of host genes with ageing across various eukaryotic hosts was investigated using the CellAge and GenAge databases (52). Podder et al. used, among other databases, the Digital Ageing Atlas to discover longevity genes associated with nutrient sensing (53). A new variant in HLA-DQB1 gene was associated with longevity and lipid homeostasis in a Chinese population study that applied LongevityMap to variant selection (54). Finally, a Cardoso et al. used several HAGR databases to generate biomarker panels for human frailty (55). These instances underscore the diverse ways in which HAGR databases can support ageing research.

## Discussion

Ageing is a complex process that arises from the interplay of various molecular pathways and the environment; therefore, the study of its multiple components is pivotal for understanding ageing and developing interventions. HAGR was conceived to facilitate such comprehensive analyses by offering comprehensive, consistent, and accurate data.

Compared to previous versions of HAGR, our resources have seen significant updates and growth. As illustrated in Figure 4A, since its inception in 2004, HAGR has consistently expanded in terms of users and citations, emphasizing its importance within the scientific community. Among our primary resources, AnAge and GenAge continue to attract the most visitors (Figure 4B).

In addition to HAGR, other websites and databases also provide valuable resources for studying ageing. One noteworthy example is the Ageing Atlas, an online resource that employs diverse data to explore the ageing process in a multidimensional way (56). Another database, AgeFactDB, compiles ageing-related factors, incorporating our databases in its analyses (57). Nonetheless, HAGR stands out as a leading resource in biogerontology due to its integrated features, offering comprehensive tools, datasets, and insights into ageing and longevity. We anticipate that HAGR, together with other tools, will continue to advance the study of ageing biology and contribute to our overarching goal: developing a paradigm that explains ageing and improves human health and longevity.

## Acknowledgements

We are grateful to past and present lab members and to the numerous individuals who have contributed our databases, with special recognition to Steve Austad for advice on AnAge. HAGR is developed active user input, and are thankful to the many users who have provided feedback and suggestions that have helped us improve our resources.

## References

1. Lopez-Otin, C., Blasco, M.A., Partridge, L., Serrano, M. and Kroemer, G. (2013) The hallmarks of aging. Cell, 153, 1194–1217.

2. Gems, D. and de Magalhaes, J.P. (2021) The hoverfly and the wasp: A critique of the hallmarks of aging as a paradigm. Ageing Res Rev, 70, 101407.

3. Rose, M.R. (2009) Adaptation, aging, and genomic information. Aging (Albany NY), 1, 444–450.

4. de Magalhaes, J.P., Costa, J. and Toussaint, O. (2005) HAGR: the Human Ageing Genomic Resources. Nucleic acids research, 33, D537–543.

5. Tacutu, R., Thornton, D., Johnson, E., Budovsky, A., Barardo, D., Craig, T., Diana, E., Lehmann, G., Toren, D., Wang, J. et al. (2018) Human Ageing Genomic Resources: new and updated databases. Nucleic Acids Res, 46, D1083–D1090.

6. de Magalhães, J.P. and Toussaint, O. (2004) GenAge: a genomic and proteomic network map of human ageing. FEBS Lett, 571, 243–247.

7. Palmer, D., Fabris, F., Doherty, A., Freitas, A.A. and de Magalhães, J.P. (2021) Ageing transcriptome meta-analysis reveals similarities and differences between key mammalian tissues. Aging (Albany NY), 13, 3313–3341.

8. de Magalhaes, J.P., Curado, J. and Church, G.M. (2009) Meta-analysis of age-related gene expression profiles identifies common signatures of aging. Bioinformatics, 25, 875–881.

9. Ohtsuka, H., Shimasaki, T. and Aiba, H. (2021) Genes affecting the extension of chronological lifespan in Schizosaccharomyces pombe (fission yeast). Mol Microbiol, 115, 623–642.

10. Lin, S.J. and Austriaco, N. (2014) Aging and cell death in the other yeasts, Schizosaccharomyces pombe and Candida albicans. FEMS Yeast Res, 14, 119–135.

11. de Magalhaes, J.P. and Costa, J. (2009) A database of vertebrate longevity records and their relation to other life-history traits. J Evol Biol, 22, 1770–1774.

12. Barardo, D., Thornton, D., Thoppil, H., Walsh, M., Sharifi, S., Ferreira, S., Anzic, A., Fernandes, M., Monteiro, P., Grum, T. et al. (2017) The DrugAge database of aging-related drugs. Aging cell, 16, 594–597.

13. Moskalev, A., Guvatova, Z., Lopes, I.A., Beckett, C.W., Kennedy, B.K., De Magalhaes, J.P. and Makarov, A.A. (2022) Targeting aging mechanisms: pharmacological perspectives. Trends Endocrinol Metab, 33, 266–280.

14. de Magalhães, J.P. (2021) Longevity pharmacology comes of age. Drug Discov Today, 26, 1559–1562.

15. Campisi, J., Kapahi, P., Lithgow, G.J., Melov, S., Newman, J.C. and Verdin, E. (2019) From discoveries in ageing research to therapeutics for healthy ageing. Nature, 571, 183–192.

16. Santos, J., Leitão-Correia, F., Sousa, M.J. and Leão, C. (2016) Dietary Restriction and Nutrient Balance in Aging. Oxid Med Cell Longev, 2016, 4010357.

17. Fontana, L., Ghezzi, L., Cross, A.H. and Piccio, L. (2021) Effects of dietary restriction on neuroinflammation in neurodegenerative diseases. J Exp Med, 218.

18. de Magalhaes, J.P., Wuttke, D., Wood, S.H., Plank, M. and Vora, C. (2012) Genome-environment interactions that modulate aging: powerful targets for drug discovery. Pharmacol Rev, 64, 88–101.

19. Wuttke, D., Connor, R., Vora, C., Craig, T., Li, Y., Wood, S., Vasieva, O., Shmookler Reis, R., Tang, F. and de Magalhaes, J.P. (2012) Dissecting the gene network of dietary restriction to identify evolutionarily conserved pathways and new functional genes. PLoS Genet, 8, e1002834.

20. Plank, M., Wuttke, D., van Dam, S., Clarke, S.A. and de Magalhaes, J.P. (2012) A meta-analysis of caloric restriction gene expression profiles to infer common signatures and regulatory mechanisms. Mol Biosyst, 8, 1339–1349.

21. Di Micco, R., Fumagalli, M., Cicalese, A., Piccinin, S., Gasparini, P., Luise, C., Schurra, C., Garre’, M., Nuciforo, P.G., Bensimon, A. et al. (2006) Oncogene-induced senescence is a DNA damage response triggered by DNA hyper-replication. Nature, 444, 638–642.

22. de Magalhaes, J.P. and Passos, J.F. (2018) Stress, cell senescence and organismal ageing. Mechanisms of ageing and development, 170, 2–9.

23. Katlinskaya, Y.V., Carbone, C.J., Yu, Q. and Fuchs, S.Y. (2015) Type 1 interferons contribute to the clearance of senescent cell. Cancer Biol Ther, 16, 1214–1219.

24. Sagiv, A. and Krizhanovsky, V. (2013) Immunosurveillance of senescent cells: the bright side of the senescence program. Biogerontology, 14, 617–628.

25. Olivieri, F., Prattichizzo, F., Grillari, J. and Balistreri, C.R. (2018) Cellular Senescence and Inflammaging in Age-Related Diseases. Mediators Inflamm, 2018, 9076485.

26. Campisi, J., Andersen, J.K., Kapahi, P. and Melov, S. (2011) Cellular senescence: a link between cancer and age-related degenerative disease? Semin Cancer Biol, 21, 354–359.

27. Palmer, A.K., Tchkonia, T., LeBrasseur, N.K., Chini, E.N., Xu, M. and Kirkland, J.L. (2015) Cellular Senescence in Type 2 Diabetes: A Therapeutic Opportunity. Diabetes, 64, 2289–2298.

28. Martínez-Cué, C. and Rueda, N. (2020) Cellular Senescence in Neurodegenerative Diseases. Front Cell Neurosci, 14, 16.

29. McCulloch, K., Litherland, G.J. and Rai, T.S. (2017) Cellular senescence in osteoarthritis pathology. Aging Cell, 16, 210–218.

30. Avelar, R.A., Ortega, J.G., Tacutu, R., Tyler, E.J., Bennett, D., Binetti, P., Budovsky, A., Chatsirisupachai, K., Johnson, E., Murray, A. et al. (2020) A multidimensional systems biology analysis of cellular senescence in aging and disease. Genome biology, 21, 91.

31. Tejada-Martinez, D., Avelar, R.A., Lopes, I., Zhang, B., Novoa, G., de Magalhães, J.P. and Trizzino, M. (2021) Positive selection and enhancer evolution shaped lifespan and body mass in great apes. Mol Biol Evol.

32. Chatsirisupachai, K., Palmer, D., Ferreira, S. and de Magalhaes, J.P. (2019) A human tissuespecific transcriptomic analysis reveals a complex relationship between aging, cancer, and cellular senescence. Aging cell, 18, e13041.

33. Brooks-Wilson, A.R. (2013) Genetics of healthy aging and longevity. Hum Genet, 132, 1323–1338.

34. Budovsky, A., Craig, T., Wang, J., Tacutu, R., Csordas, A., Lourenco, J., Fraifeld, V.E. and de Magalhaes, J.P. (2013) LongevityMap: a database of human genetic variants associated with longevity. Trends Genet, 29, 559–560.

35. Craig, T., Smelick, C., Tacutu, R., Wuttke, D., Wood, S.H., Stanley, H., Janssens, G., Savitskaya, E., Moskalev, A., Arking, R. et al. (2015) The Digital Ageing Atlas: integrating the diversity of agerelated changes into a unified resource. Nucleic acids research, 43, D873–878.

36. Keane, M., Craig, T., Alfoldi, J., Berlin, A.M., Johnson, J., Seluanov, A., Gorbunova, V., Di Palma, F., Lindblad-Toh, K., Church, G.M. et al. (2014) The Naked Mole Rat Genome Resource: facilitating analyses of cancer and longevity-related adaptations. Bioinformatics, 30, 3558–3560.

37. Keane, M., Semeiks, J., Webb, A.E., Li, Y.I., Quesada, V., Craig, T., Madsen, L.B., van Dam, S., Brawand, D., Marques, P.I. et al. (2015) Insights into the evolution of longevity from the bowhead whale genome. Cell reports, 10, 112–122.

38. de Magalhaes, J.P., Cabral, J.A. and Magalhaes, D. (2005) The influence of genes on the aging process of mice: a statistical assessment of the genetics of aging. Genetics, 169, 265–274.

39. Berkel, C. and Cacan, E. (2021) Analysis of longevity in Chordata identifies species with exceptional longevity among taxa and points to the evolution of longer lifespans. Biogerontology, 22, 329–343.

40. Whittemore, K., Vera, E., Martinez-Nevado, E., Sanpera, C. and Blasco, M.A. (2019) Telomere shortening rate predicts species life span. Proceedings of the National Academy of Sciences of the United States of America, 116, 15122–15127.

41. Kerepesi, C., Meer, M.V., Ablaeva, J., Amoroso, V.G., Lee, S.G., Zhang, B., Gerashchenko, M.V., Trapp, A., Yim, S.H., Lu, A.T. et al. (2022) Epigenetic aging of the demographically non-aging naked mole-rat. Nat Commun, 13, 355.

42. Castiglione, G.M., Zhou, L., Xu, Z., Neiman, Z., Hung, C.F. and Duh, E.J. (2021) Evolutionary pathways to SARS-CoV-2 resistance are opened and closed by epistasis acting on ACE2. PLoS Biol, 19, e3001510.

43. Cagan, A., Baez-Ortega, A., Brzozowska, N., Abascal, F., Coorens, T.H.H., Sanders, M.A., Lawson, A.R.J., Harvey, L.M.R., Bhosle, S., Jones, D. et al. (2022) Somatic mutation rates scale with lifespan across mammals. Nature, 604, 517–524.

44. Kitazoe, Y., Kishino, H., Tanisawa, K., Udaka, K. and Tanaka, M. (2019) Renormalized basal metabolic rate describes the human aging process and longevity. Aging Cell, 18, e12968.

45. Townes, F.W., Carr, K. and Miller, J.W. (2020) Identifying longevity associated genes by integrating gene expression and curated annotations. PLoS Comput Biol, 16, e1008429.

46. Dönertaş, H.M., Fuentealba, M., Partridge, L. and Thornton, J.M. (2019) Identifying Potential Ageing-Modulating Drugs In Silico. Trends Endocrinol Metab, 30, 118–131.

47. Vega Magdaleno, G.D., Bespalov, V., Zheng, Y., Freitas, A.A. and de Magalhaes, J.P. (2022) Machine learning-based predictions of dietary restriction associations across ageing-related genes. BMC Bioinformatics, 23, 10.

48. Berkel, C. and Cacan, E. (2021) A collective analysis of lifespan-extending compounds in diverse model organisms, and of species whose lifespan can be extended the most by the application of compounds. Biogerontology, 22, 639–653.

49. Xu, P., Wang, M., Song, W.M., Wang, Q., Yuan, G.C., Sudmant, P.H., Zare, H., Tu, Z., Orr, M.E. and Zhang, B. (2022) The landscape of human tissue and cell type specific expression and coregulation of senescence genes. Mol Neurodegener, 17, 5.

50. Choi, I., Wang, M., Yoo, S., Xu, P., Seegobin, S.P., Li, X., Han, X., Wang, Q., Peng, J., Zhang, B. et al. (2023) Autophagy enables microglia to engage amyloid plaques and prevents microglial senescence. Nat Cell Biol, 25, 963–974.

51. Neumann, B., Baror, R., Zhao, C., Segel, M., Dietmann, S., Rawji, K.S., Foerster, S., McClain, C.R., Chalut, K., van Wijngaarden, P. et al. (2019) Metformin Restores CNS Remyelination Capacity by Rejuvenating Aged Stem Cells. Cell Stem Cell, 25, 473-485.e478.

52. Teulière, J., Bernard, C. and Bapteste, E. (2021) Interspecific interactions that affect ageing: Age-distorters manipulate host ageing to their own evolutionary benefits. Ageing Res Rev, 70, 101375.

53. Podder, A., Raju, A. and Schork, N.J. (2021) Cross-Species and Human Inter-Tissue Network Analysis of Genes Implicated in Longevity and Aging Reveal Strong Support for Nutrient Sensing. Front Genet, 12, 719713.

54. Yang, F., Sun, L., Zhu, X., Han, J., Zeng, Y., Nie, C., Yuan, H., Li, X., Shi, X., Yang, Y. et al. (2017) Identification of new genetic variants of HLA-DQB1 associated with human longevity and lipid homeostasis-a cross-sectional study in a Chinese population. Aging (Albany NY), 9, 2316–2333.

55. Cardoso, A.L., Fernandes, A., Aguilar-Pimentel, J.A., de Angelis, M.H., Guedes, J.R., Brito, M.A., Ortolano, S., Pani, G., Athanasopoulou, S., Gonos, E.S. et al. (2018) Towards frailty biomarkers: Candidates from genes and pathways regulated in aging and age-related diseases. Ageing Res Rev, 47, 214–277.

56. Consortium, A.A. (2021) Aging Atlas: a multi-omics database for aging biology. Nucleic Acids Res, 49, D825–D830.

57. Hühne, R., Thalheim, T. and Sühnel, J. (2014) AgeFactDB--the JenAge Ageing Factor Database--towards data integration in ageing research. Nucleic Acids Res, 42, D892–896.

